# Career Choice, Gender, and Mentor Impact: Results of the U.S. National Postdoc Survey

**DOI:** 10.1101/355511

**Authors:** Sean C. McConnell, Erica L. Westerman, Joseph F. Pierre, Erin J. Heckler, Nancy B. Schwartz

## Abstract

The postdoctoral community is an essential component of the academic and scientific workforce. As economic and political pressures impacting these enterprises continue to change, the postdoc experience has evolved from short, focused periods of training into often multidisciplinary, extended positions with less clear outcomes. As efforts are underway to amend U.S. federally funded research policies, the paucity of postdoc data has made evaluating the impact of policy recommendations challenging. Here we present comprehensive survey results from over 7,600 postdocs based at 351 academic and non-academic U.S. institutions in 2016. In addition to demographic and salary information, we present multivariate analyses on the factors that influence postdoc career plans and mentorship satisfaction in this population. We further analyze gender dynamics and expose wage disparities and career choice differences. Academic research positions remain the predominant career choice of postdocs in the U.S., although unequally between postdocs based on gender and residency status. Receiving mentorship training during the postdoctoral period has a large, positive effect on postdoc mentorship satisfaction. Strikingly, the quality of and satisfaction with postdoc mentorship appears to also heavily influence career choice. The data presented here are the most comprehensive data on the U.S. postdoc population to date. These results provide an evidence basis for informing government and institutional policies, and establish a critical cornerstone for quantifying the effects of future legislation aimed at the academic and scientific workforce.

## Introduction

Postdoctoral training offers doctoral recipients a temporary period of mentored or scholarly experience, considered highly productive within scientific and academic communities. Such training is also ostensibly valuable for postdocs, who gain additional experience to help pursue their chosen career paths. Tenure-track faculty positions, however, are now estimated to represent a small percentage of postdoc career outcomes (~15%)^1,2^. This has led to proposals to support training postdocs for additional roles beyond tenure-track faculty positions, and additional efforts by the National Institutes of Health (NIH), National Science Foundation (NSF), and National Academies of Sciences, Engineering, and Medicine to increase mentor accountability^2–6^. Persistent concerns with increasingly long periods of postdoctoral training, lack of appropriate career guidance beyond the professoriate, and comparatively low postdoctoral salaries, have also led to repeated calls to reform the postdoctoral training model^7–12^. Despite these concerns, comprehensive data for postdocs are not routinely collected^2,4^. Indeed, reliable data on such basic information as the number of postdocs have been lacking, or disputed, in part due to difficulties in collecting these data because of lack of job title standardization, postdoc mobility, and the ad hoc nature of institutional postdoctoral administration^2,5,12–14^.

Possibly for these very reasons, the postdoctoral experience has not been comprehensively surveyed nationally in over a decade, following the Sigma Xi “Doctors without Orders” survey report in 2005, which was based on postdoc respondents from 46 participating institutions^8^. Nevertheless, recent data collection efforts have provided insights into the postdoctoral experience^5,8,9,14–19^. For example, the pilot phase of the NSF Early Career Doctorates Survey studied the breadth of the doctoral population at U.S. academic institutions, including postdoctoral researchers (31%), young faculty (54%), and scientists in non-postdoctoral positions^18^. Yet, to date comprehensive survey data specifically targeting the postdoctoral period and including postdoctoral researchers with PhDs granted both inside and outside the U.S., as well as data regarding postdoc career plans, mentor satisfaction, and demographics are still largely lacking^2,4^. To address these gaps, and to research those without clear institutional oversight, we took a grass-roots approach to conduct a postdoc-led survey of U.S. postdoctoral researchers. We asked postdoctoral researchers a number of questions associated with professional and career development, mentoring, career choice, lifestyle, and demographics (for details see Materials and Methods and Data S1-S3). The purpose of this work was to capture a comprehensive snapshot of the postdoctoral experience in a manner that was both broad and informative, with high diversity in questions and topics covered, in the number of institutions, and in the breadth of postdoctoral experiences included.

## Results and Discussion

To collect data from institutions with a wide range of support for postdocs, we took a multi-level approach to recruit survey participants. We used publicly available contact information for university leadership, postdoctoral administrators, postdoctoral societies and associations, and asked these people in leadership positions to disseminate our survey to all postdocs at their institutions. In total, we contacted individuals at 482 institutions most likely to have postdocs, including universities, research institutions, museums, and government labs. We obtained respondents from 351 institutions. In addition to direct contact with institutions, we also used a grass-roots survey dissemination approach, promoted a website describing the survey that could be freely shared on social media and by email, and contacted professional societies to encourage survey dissemination. Using these combined approaches, we collected 7,673 individual responses into a secure REDCap database (IRB Protocol Number 15-1724), which, after quality control to remove respondents from non-U.S. institutions, provided a final dataset of 7,603 respondents (see Materials and Methods for details). As one of our goals was to reach as many postdocs as possible, survey dissemination was not randomized to any specific subset. Responses from institutions with long-standing postdoctoral affairs offices were anticipated to be over-represented in our dataset (see Materials and Methods for more information). Nevertheless, our respondents represented all 50 states, including a large fraction of respondents from institutions without well-established offices for postdoctoral support. While the majority of respondents represented STEM disciplines, which traditionally employ the most postdocs, 8.4% reported their primary fields as humanities, psychology, or social sciences (Table S3).

Our postdoc respondents were 49% U.S. citizens and 51% non-U.S. citizens (Fig. 1 and Table S2). The majority were 30-34 years of age (54.5%), and 1-3 years from receipt of their doctorate (63.1%), matching their reported years of postdoctoral experience (Fig. 1 and Table S2). The majority (55%) described their primary field of study as life sciences. There were small, but significant, differences in primary field by geographic region (Fig. 2 and Table S3). Race and ethnicity were self-reported with 60.3% White/Caucasian, 24.8% Asian/Asian American, 6.6% Hispanic/Latino, and 2.6% Black/African American (Table S2). Both national and international postdocs were included in these proportions. Our respondents were 53% female, while the gender ratio of their mentors was skewed towards males (71% male; Fig. 1), consistent with the most recent AAUP Gender Equity report where full-time faculty are majority (61%) male^20^.

**Figure 1:**
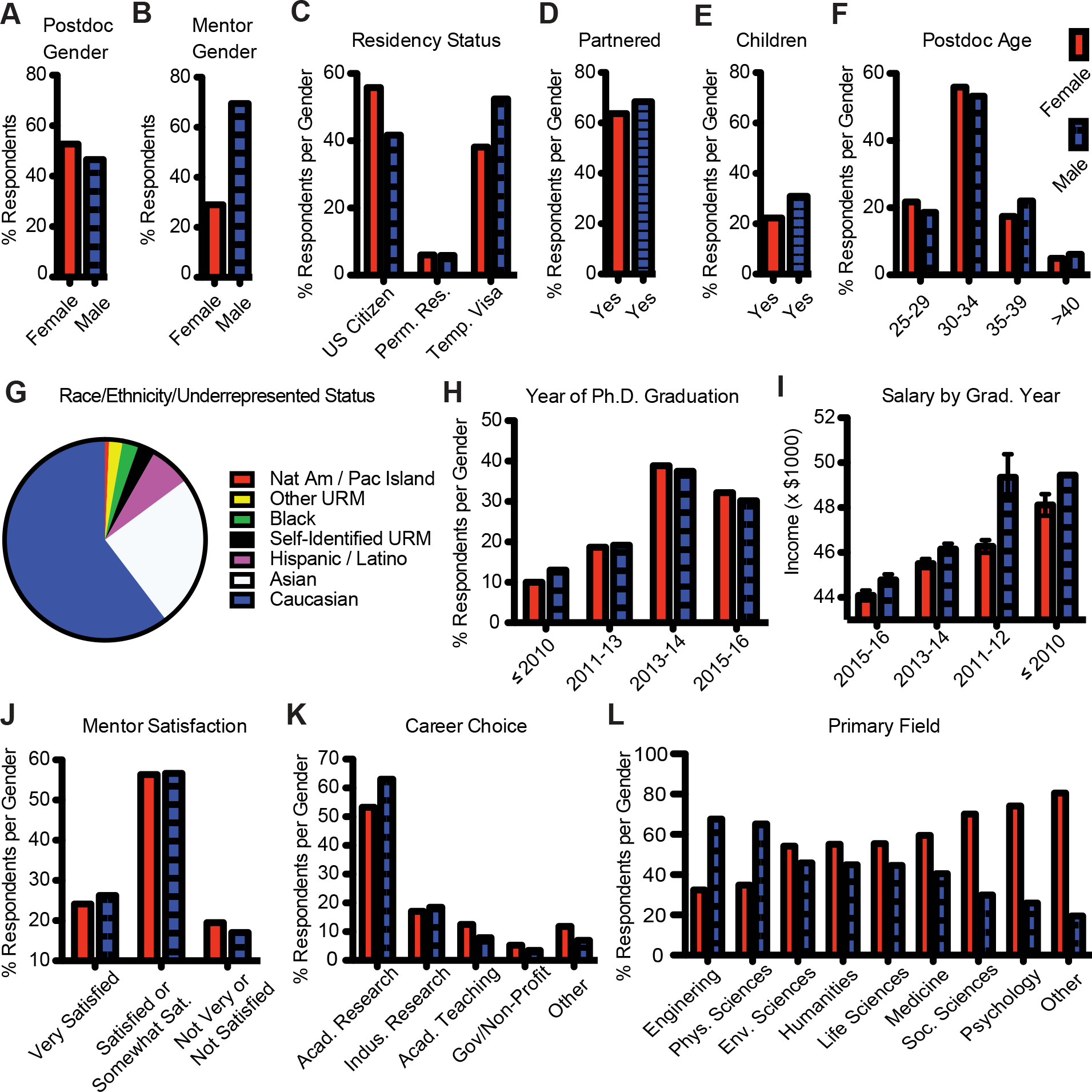
Demographics of the Postdoc Population Surveyed. A) Postdoc gender, B) Mentor gender, C) Residency status, D) Partnered/Married, E) Has children, F) Age, G) Race/Ethnicity/Underrepresented status (which may include things other than race and ethnicity), H) Year of graduation, I) Adjusted income, by year of graduation, J) Postdoc satisfaction with mentor, K) Primary long-term career plans, and L) Primary field/discipline. Red bars indicate female, striped blue bars indicate male.

**Figure 2:**
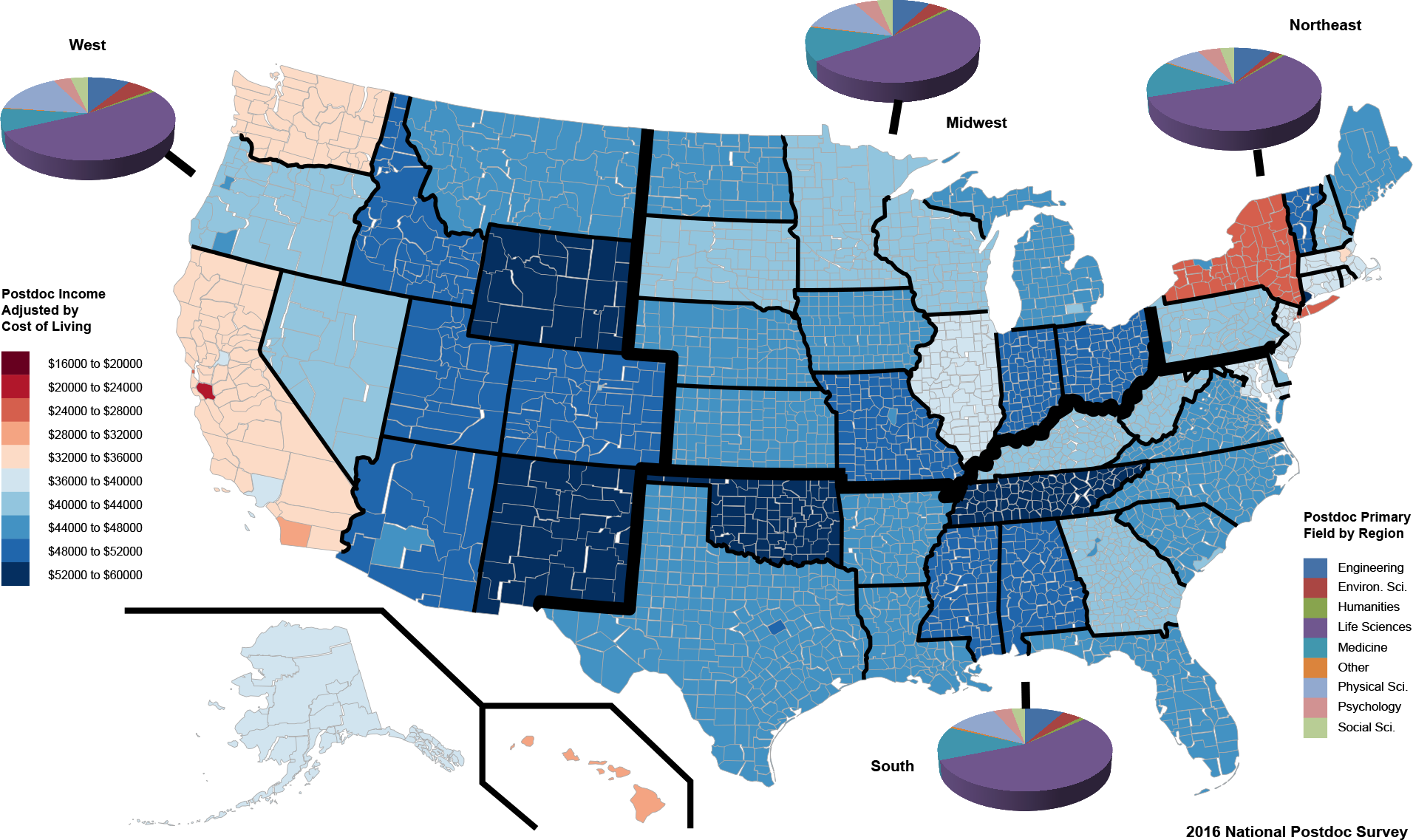
Postdoc Cost of Living Adjusted Income and Field of Study by Region. A map of the United States with the range of reported postdoc gross income adjusted by cost of living (key on the left) and the respondents’ field of study (key on the right) in each of the four major regions: West, Midwest, South, and Northeast (bold lines). The adjusted income data are provided at the state (and when data sufficient to support, county) level.

While the demographics of our survey respondents may differ slightly from those of the actual postdoctoral population (but see Materials and Methods for analysis suggesting lack of response bias), confirmation of a lack of response bias remains difficult as there are currently no gold-standard datasets of postdocs in the U.S. for comparison, due to the previously mentioned broader lack of oversight and barriers to reaching postdocs. That being said, the unique characteristics of our dataset, including approximately equal representation (similar sample sizes) of men and women, as well as U.S. citizens and non-U.S. citizens, facilitated our comparative analyses of the postdoctoral experiences of these different groups, which we report below.

Our data indicate that gender has a significant effect on the postdoc experience (Fig. 1). Men were paid more than women (Male average: $47,678.00, Female average: $46, 477.43, n=7,516, χ^2^=62.337, p<0.0001). Men were more likely to have a same-gender mentor (i.e. a same-gender role model) (Male: 77.3%, Female: 35.4%, n=7,459, χ^2^=144.352, p<0.0001). Men were more likely to be non-U.S. citizens (Male: 42% U.S. citizens, 52% temporary visas, 6% permanent residents; Female: 56% U.S. citizens, 38% temporary visas, 6% permanent residents; n=7,543, χ^2^=169.709, p<0.0001). In addition, a small but significantly higher proportion of male postdocs were married/partnered (Male: 68.3%, Female: 63.2%, n=7,538, χ^2^=21.693, p<0.0001) and/or have children (Male: 31.0%, Female: 22.3%, n=7,532, χ^2^=71.561, p<0.0001). The gender disparity in pay occurs even after male and female postdocs were matched in age, years since graduation, mentor satisfaction, and likelihood of being married/partnered (nominal logistic model, gender effect test n=7,311, χ^2^=47.235, p<0.0001, Table S4). This gender wage gap increased with postdoc age but not with partnership status, partially supporting previous analyses of the STEM gender wage gap^20,21^. Male postdocs were also more likely than women to have received PhD degrees in Engineering (n=620, χ^2^=76.652, p<0.0001) or the Physical Sciences (n=846, χ^2^=77.466, p<0.0001), two fields which have historically higher salaries^22^. Interestingly, female postdocs trended towards being paid less than men in all fields except the Physical Sciences, where women trended towards being paid slightly more than men (Table S5). Income, mentor gender, citizenship, and partner status are all factors that may contribute to the observed gender difference in interest in primarily research-focused academic careers^22^ (Fig. 1 & 3G).

Most postdocs reported salaries in the range of $39,000 − $55,000 (median $43,750; mean $46,988, n=7,551). In the 2014 National Postdoctoral Association’s Institutional Policy Report, 52% of the 74 institutions reported their minimum stipend matched the current NIH NRSA minimum^17^. At the time of this survey, the NIH minimum^23^ was $43,692, which matches well with $43,750, the reported median income in our study. Five percent of postdocs reported mean gross incomes of less than $39,000 and ~10% reported incomes above $55,000. Although salaries in high cost of living (COL) urban areas tend to be higher than average (Table S6), when adjusted to publicly available COL data, postdocs in large metropolitan areas earn significantly less money than postdocs in college towns or rural settings (average salary when adjusted for cost of living, metropolitan: $38,045.60, non-metropolitan: $44,714.40; n=7,551, F-ratio: 12.614, p=0.0002) (Fig. 2, Table S6). “The Postdoctoral Experience Revisited” 2014 report recommended as a best practice that the minimum salary be set at $50,000; however, this has not been enacted at most institutions or by funding agencies^3^. During the months that our survey was open (February — September 2016), the effect of a proposed minimum salary update ($47,476) to the Fair Labor Standards Act (FLSA) on postdoctoral salaries was openly debated, but ultimately not federally mandated^24^. Our data suggest that setting a minimum salary for postdocs is particularly important for postdoctoral researchers in large metropolitan areas, where salaries are not maintaining parity with cost of living increases.

The majority of postdocs indicated research-focused academic careers as their primary long term career plan (57.7%), with industry research a distant second (17.8%, Fig. 1J and Table S2). Determining the “why” of career choice remains the subject of much study^2,4,5^. To assess which factors were most influential for determining postdoc career plan in our dataset, (categorized in this survey as: academia, primarily research based; academia, primarily teaching based; industry; government/non-profit; other) we conducted a nominal logistical model with 26 factors concerning topics considered to be important for postdoc success and career choice (Table S7), which include demographics, training, productivity and mentor support matrices. The 14 significant factors in the model were (in order of effect size): 1) whether postdoc career plans had changed; 2) whether training in pedagogy was received; 3) feelings of career preparedness; 4) perceived mentor support of career plan; 5) primary field of study; 6) residency status in the U.S.; 7) intensity of job search; 8) postdoc gender; 9) number of first, last, or corresponding author publications; 10) number of conferences attended in the past year; 11) hours worked per week; 12) total number of publications while a postdoc; 13) mentor rank; and 14) desire to pursue a career in the U.S. (Table 1). Perceived mentor support, number of postdoc publications, hours worked per week, conferences attended, and postdoc feelings of career preparedness were all positively correlated with choice to pursue a research-focused academic career (Fig. 3A, C-F). Male postdocs, and postdocs who were not U.S. citizens, were more interested in academic research positions (Fig. 3G & 3H). In contrast, postdocs with mentors outside of the professoriate were more likely to prefer government/non-profit positions (Fig. 3B). Whether this is a cause or effect relationship is not clear in our study, though we did find that postdocs with non-academic mentors changed their career plans at the same rate as those with academic mentors (n=7,361, χ^2^=6.860, p=0.077). In addition, postdocs actively searching for permanent positions were less interested in academic research than postdocs not yet on the job market (Fig. 3I), and were more likely to have changed their career plans (n=7,565, χ^2^=224.633, p<0.0001). These results complement recent studies suggesting that individual career choice is influenced by changing job attribute preferences and self-awareness^22^, and that academic success is influenced by mentorship during the postdoctoral period^26^.

**Table 1.** Significant factors influencing postdoc primary career plans. Whole model effect: n=6,504, Model R^2^=0.2017, AICc=15924, BIC=21130.

| Factor | $\chi^2$ | -log p-value |
| --- | --- | --- |
| Whether long term career plans have changed | 599.951 | 108.529 |
| Received training in pedagogy | 151.052 | 27.273 |
| Feelings of career preparedness | 161.510 | 11.925 |
| Perceived mentor support of career plan | 130.577 | 11.925 |
| Primary field of study | 191.331 | 10.190 |
| Residency status in U.S. | 133.264 | 9.941 |
| Job search intensity | 98.574 | 9.352 |
| Postdoc gender | 53.654 | 7.658 |
| Number of first, last, or corresponding author publications | 86.193 | 5.274 |
| Conferences attended in last year | 84.468 | 5.043 |
| Hours worked / week | 109.093 | 4.870 |
| Total number of publications while a postdoc | 80.503 | 4.524 |
| Academic rank of mentor | 70.513 | 3.292 |
| Plan to pursue a career in U.S. | 37.452 | 2.340 |

**Figure 3:**
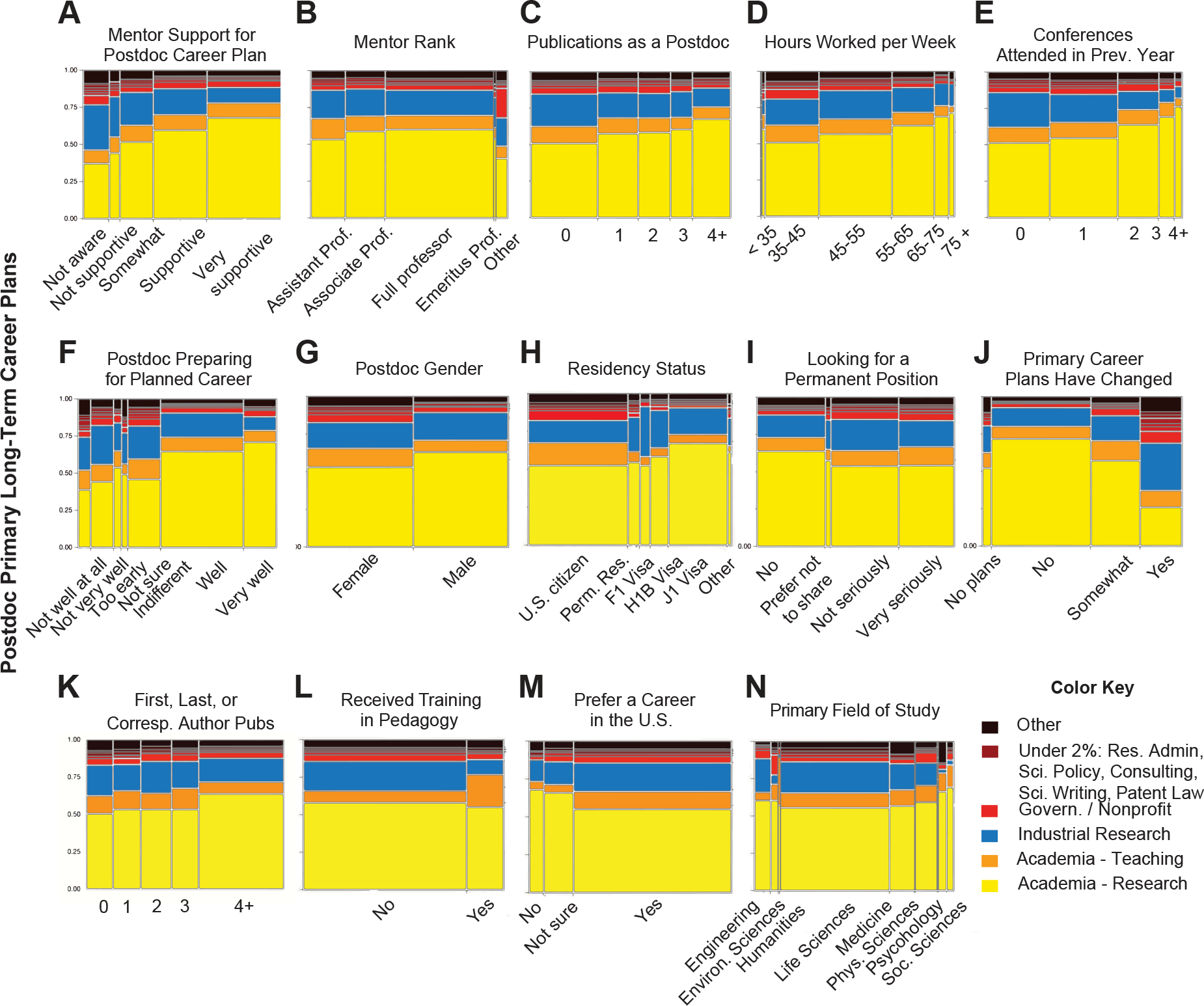
Postdoc Career Choice. Here we illustrate the independent effects of the fourteen significant factors (out of twenty-six) in the nominal logistic regression model of best fit for postdoc primary career choice. In these mosaic plots, the panels show the listed factor and corresponding effect size, and the right-hand color key corresponds to primary career choice. Factors are paraphrased survey questions, please see Data S1 and S2 for specific wording of questions.

Sixty percent of respondents were satisfied with the mentorship they receive, with similar responses from both genders (Fig. 1I). To assess which factors were most influential for determining postdoc mentor satisfaction, we conducted a nominal logistic model with the same 26 factors included in the model for postdoc career choice (though excluding mentor satisfaction as a factor, and replacing it with postdoc long term career plan) (Table S7). The 8 significant factors in the model (in order of effect size) were: 1) feelings of career preparedness; 2) perceived mentor support of career plan; 3) frequency of project meetings with mentor; 4) intensity of job search; 5) whether training in mentorship was received; 6) primary field of study; 7) perception of job market; and 8) academic rank of mentor (Table 2). These factors were more important than number of postdoc publications, whether a postdoc had changed career plans, postdoc or mentor gender, residency status, or training in either grant writing or pedagogy.

**Table 2.** Significant factors influencing postdoc satisfaction with mentoring. Whole model effect: n=6,504, Model R^2^=0.3007, AICc=14729, BICc=17810.

| Factor | $\chi^2$ | -log p-value |
| --- | --- | --- |
| Feelings of career preparedness | 960.457 | 181.948 |
| Perceived mentor support of career plan | 904.891 | 178.146 |
| Frequency of mentor meetings | 532.31 | 89.480 |
| Job search intensity | 68.255 | 8.040 |
| Received training in mentorship | 37.088 | 6.240 |
| Primary field of study | 92.193 | 4.368 |
| Perception of academic job market | 48.088 | 3.384 |
| Academic rank of mentor | 41.614 | 2.508 |

Perceived mentor support had a positive effect on mentor satisfaction, as did frequency of mentor meetings, perception of preparedness for desired future career, and perception of job market (Fig. 4A-D). Receiving mentor training also had a large positive effect on postdoc mentor satisfaction (Fig. 4E). We found this to be particularly noteworthy, as mentorship training is not a common part of the postdoctoral experience, with only 26% of postdocs reporting that they have received such training.

**Figure 4:**
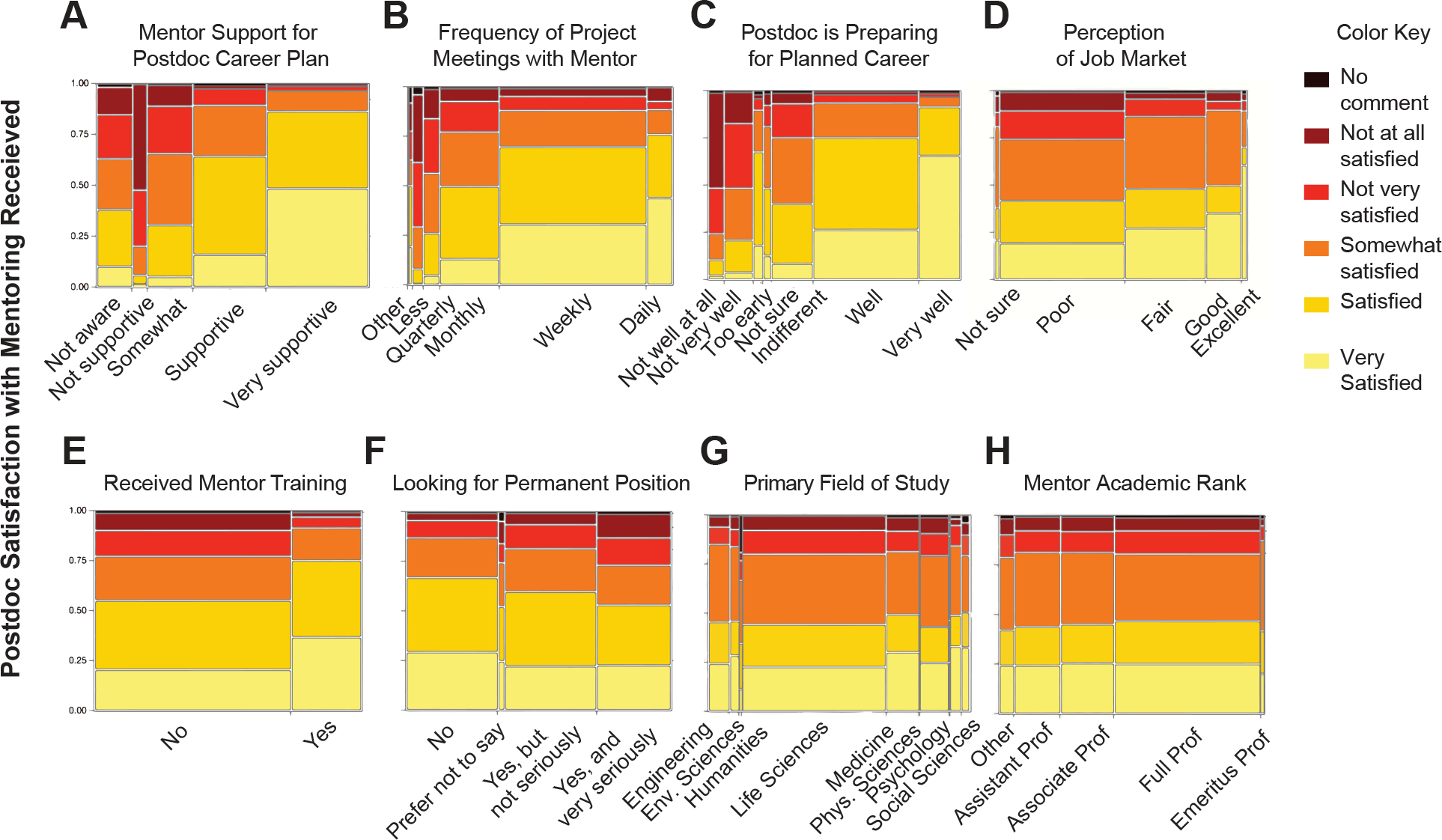
Postdoc Mentor Satisfaction. Here we illustrate the independent effects of the eight significant factors (out of twenty-six) in the nominal logistic regression model of best fit for postdoc mentor satisfaction. In these mosaic plots, the panels show the listed factor and corresponding effect size, and the right-hand color key corresponds to the degree of mentor satisfaction. Factors are paraphrased survey questions, please see Data S1 and S2 for specific wording of questions.

Previous research on a postdoc cohort showed that high mentorship satisfaction and perceived support correlated with increased interest in an academic research focused career^27^. In addition, in a randomized, controlled study, a different type of mentoring, “group career coaching,” was found to increase both perceived “achievability” and “desirability” of academic careers in an under-represented minority student group^28^. While our data do not show a significant correlation between gender and mentor satisfaction, they do suggest that an increase in mentor support and mentorship training may increase female and under-represented postdocs’ pursuit of research-intensive academic careers.

## Conclusions

In summary, our dataset represents the most comprehensive survey of the U.S. postdoctoral population in over a decade. As such, these data may provide a benchmark for legislation and institutional policy makers, inform research questions pertaining to the evolving postdoctoral population, and serve as a precedent for understanding the important dynamics of the scientific workforce. We found that a research-focused academic position remains the most common primary career goal for postdocs, in spite of increasing emphasis on other types of careers for doctorate holders^4,10,29^. Although sixty percent of respondents were satisfied with the mentoring that they receive, our data suggest that inclusion of formal mentorship training for postdocs may significantly increase mentor satisfaction and influence career choice^28^. Our data also show that women are less interested in research-focused academic positions than men, and this may be associated with gender specific differences in postdoctoral experiences^30^.

While the data we collected allowed us to identify a number of factors influencing the postdoctoral experience, other factors, such as socioeconomic background and underrepresented status may also play a significant role, and should be studied further. Nevertheless, our findings highlight the impact of mentoring, across all demographics, as essential to informing career choice and determining quality of postdoctoral experience.

## Materials and Methods

### Survey instrument design

The National Postdoc Survey questions were designed to emphasize aspects of the postdoctoral experience related to career choice and mentoring, in addition to collection of basic demographics. These questions were based on over a decade of experience with postdoctoral surveys administered at the University of Chicago, led by postdocs within the Biological Sciences Division Postdoctoral Association. In an effort to maximize participation for all postdocs, regardless of institutional environment, we disseminated the survey using top-down and grassroots methods described below.

We conducted the survey in two phases, a 15-instutution pilot phase followed by a national rollout to over 450 institutions. The pilot phase was launched on February 2, 2016 after contacting and inviting participation from administrators at the 15 member schools of the Committee on Institutional Cooperation (CIC, now the Big Ten Academic Alliance plus the University of Chicago). 272 postdocs participated in the pilot phase of the survey. Feedback about the survey design was solicited during a workshop about the survey presented at the National Postdoc Association Annual meeting on March 4, 2016. The pilot survey questions (Data S1) were then slightly modified before nationwide launch on March 31, 2016. These revisions included additional demographic questions, and rephrasing of several questions to improve clarity (Data S2 and S3). The revised survey was available from March 31-September 2, 2016. While the CIC institutions participated in the pilot version of this survey, the survey was also open to postdocs at CIC institutions after the national rollout. A majority of participants from CIC institutions responded after March 31, 2016 and took the final version of the survey rather than the pilot version.

For the top-down survey dissemination approach, a team of five postdocs and two administrators compiled contacts for all doctoral degree and research institutions in the U.S. that were thought to have postdoctoral researchers. We gathered publically available contact information for Postdoctoral Offices, Postdoctoral Associations, as well as Offices of Research, Deans of Graduate Schools, Provosts and any other administrators that may represent postdocs for each institution (including website, email addresses, names) via web search. Whenever an institution did not have a postdoctoral office, we tried to determine who had oversight over postdoctoral researchers such as a representative from an Office of Research, Graduate School, or a Provost Office. We used this information to simultaneously contact those who we determined were most likely to represent postdocs at each University, including any listed postdoc contacts. Multi-respondent emails were sent to the above described representatives at each institution. These individuals were again invited to participate during the months of April, June, July, and August, and contact lists were revised to update contact information, and include additional institutions expected to have postdocs.

For our grass-roots survey dissemination approach we launched a website that could be freely shared on social media and by email, which explained the survey aims and contained a centralized contact form. The contact form allowed any postdocs who had not been reached via standard institutional contacts to participate in the survey through this secondary means of contact. In addition, we periodically checked contact information for institutional representatives, and updated the contact information, added new institutional contacts, and encouraged grass-roots survey dissemination during the seven months that the survey was active.

In all, 482 sets of putative postdoctoral oversight representatives were contacted by email, although some larger institutions such as Harvard University and NIH often housed separate institutes or offices that were each contacted separately - in these cases 5 and 30 sets, respectively. During the seven months (February 2-September 2, 2016) that the survey was open, over 7,600 postdoc responses were collected, with respondents from every state, and from 351 institutions and universities. While the number of respondents varied between months (ranging from 24 during the two days the survey was open in September to 2,268 in August), there was no statistical difference in the gender ratio of respondents over the seven months (whole dataset: 53.1% female and 46.9% male ±5, n=7,579, χ^2^=10.703, p=0.1521; excluding non-U.S. postdocs: 53.1% female and 46.9% male ±5; F:M, n=7,560, χ^2^=10.866, p=0.1446). Respondents from the 46 institutions that participated in the 2005 Sigma Xi survey, representing institutions with long-standing institutional support for postdocs, contributed 3,126 responses, slightly less than half of all respondents. This indicates that our addition of a grassroots approach to survey dissemination contributed to a broader sampling of postdocs across different institutional environments, providing an even more comprehensive assessment of U.S. postdoctoral experiences.

Four institution classifications were added as fixed variables to the final dataset: institution classification as public or private, Carnegie classification, U.S. Census Region, and participation in the 2005 Sigma Xi Postdoctoral Survey. City and state of each institution were also added.

### Statistical analysis

Raw response data were quality-filtered to select for U.S.-based institutions and individuals who were currently in self-described postdoctoral positions. Of the 7,673 total respondents, 70 were removed from the initial dataset using these quality filters, yielding a final dataset of 7,603 U.S. postdoctoral respondents. The demographics data shown (Fig. 1) were calculated by first sorting by gender, and then sorting by the demographic of interest displayed as total percentage of respondents per gender (all panels except Fig. 1H) or by a mean +/− standard deviation (Fig. 1H) using Prism7 (GraphPad). The effect of gender on salary, having a same sex mentor, residency status, partner status, and having children was tested using a Pearson χ^2^ test, n= 7,516, 7,459, 7,543, 7,538, 7,532 respectively). Sample sizes differed because respondents were allowed to skip questions, and are therefore reported as “n” here and throughout. However, most respondents answered most survey questions, as can been seen by the similar sample sizes for these different survey questions. The effects of gender, age, years since graduation, mentor satisfaction, and likelihood of being partnered on postdoc salary were tested using a nominal logistic model, n=7,311. The effect of gender on being in the fields of engineering or the physical sciences was tested using a Pearson χ^2^ test, engineering n=620, physical sciences n=846. We used a Bonferroni correction to account for multiple testing, yielding a significance threshold of p=0.006. All statistical tests were two-sided. Statistics were performed using JMP 13.1 by SASS.

To determine what factors were significantly correlated with postdoctoral career choice and mentor satisfaction, we ran a nominal logistic model using 26 different fixed variables listed in Table S7 using the JMP 13.1 by SASS fit model platform. We then determined which factors were significant variables after controlling for multiple testing. These estimates of effect size are reported in Table 1 and Table 2. A total of 6,504 respondents answered all 26 of the questions included in this analysis.

### Cost of living and postdoc salaries

Cost of living index (COL) data for 2016 was produced by the Council for Community and Economic Research (https://www.c2er.org/). State COL data were generated by averaging across all cities that have 2016 C2ER cost of living data provided per state. Average postdoc salary from all survey respondents for each state was divided by these state COL values to produce postdoc salaries adjusted by cost of living. Whenever income was not specified, the midpoint of income range selected by the respondent was used. These values were mapped to each state with red to blue corresponding to lowest to highest adjusted salary, respectively. In addition, counties with institutions having at least 50 respondents were then mapped separately, to map adjusted postdoc salary in 38 counties with additional COL data, against the background of the state COL data, in 50 states plus Washington D.C.

### Population proportion analysis

To determine the number of individual responses required from a total population of 100,000 for 95% and 99% confidence levels, at a 5% margin of error, assuming the true population proportion being measured is between 3-50% of the total population, we conducted a population proportion analysis using the equation and definitions as described in Tintle et al ^31^ and at Select Statistical Services Limited^32^. Results are reported in Table S1.

### Data analysis of survey respondent proportions

Inconsistent definitions across institutions and lack of existing institutional contact lists for postdocs, particularly for those without postdoctoral offices and other support, can make collecting representative data for postdocs challenging^12^. Thus demographics of respondents may differ across surveys, and the postdoctoral demographics of previous survey datasets may differ from those observed in our study. To further assess our demographic data, we conducted the comparisons described below.

We compared our demographics to that of the 2005 Sigma Xi survey which is perhaps the most comparable effort to our own, having 7600 postdoc respondents, both citizens and noncitizens^8^. The 2005 Sigma Xi dataset had 42% female postdocs (51% female for U.S. citizens, and 35% female for internationals), and overall 46% U.S. citizen and permanent resident postdocs (54% temp visas). Our current survey dataset contains a higher percentage of both female postdocs (53% female) and U.S. citizen and permanent resident postdocs (55%) relative to the Sigma Xi survey from a decade ago, which may in part reflect changing demographics of the US postdoctoral population, as well difference in institutions sampled. However, the relative difference in proportion of females for U.S. and non-U.S. citizens remains consistent (approximately 15%); our U.S. citizen respondents were 60% female, while our international respondents were 46% female.

An alternative explanation for this increase in female respondents in our dataset relative to the earlier Sigma Xi survey is that women may have disproportionately responded to our survey. We tested this hypothesis by checking the University of Chicago female and male response ratio against the actual sex ratio of female and male postdocs in the Biological Sciences. Our survey respondents from the University of Chicago were 49.3% female and 50.7% male, while the actual sex ratio of female and male postdocs in the University of Chicago Biological Sciences was 46.5% female and 53.5% male, which puts our survey respondent ratio well within the standard 5% margin of error. While it is unclear how representative University of Chicago postdocs are of the national postdoctoral population, it is important to remember that the surveyed population, by definition, all have advanced degrees, work at research institutions, and are all highly likely to have strong command of the English language, even if it is not their first language. Doctorate recipients make up 2% of the U.S. national population^33^. As doctorates are a small percentage of the national population, they are likely to make up a small percentage of respondents to general national surveys. Thus response biases of surveys targeting this population may differ from those targeting the general population.

**Supplementary Material** has been uploaded for this manuscript.

## Acknowledgements

We thank Heather Titley and Giorgio Grasselli for assistance with survey instrument design and dissemination; Laurie Risner for assistance with the Big Ten Pilot Phase; and Dylan Meyer for assistance with data clean-up. We express deep gratitude to all postdocs who participated in this survey, as well as to the postdoctoral associations, administrators, and many others who helped disseminate our survey.

## Author contributions

SCM, ELW, EJH, and JFP designed the survey and analyzed data. ELW performed multivariate analysis and models. SCM, ELW, JFP, and EJH disseminated the survey. All authors contributed to writing and editing the manuscript.

## Data and materials availability

Non-privileged data used in this study are available in supplemental tables. Due to their sensitive nature, much of the raw data are privileged to prevent individual identification, in accordance with IRB protocol.

